# Alterations in Lipid Saturation Trigger Remodeling of the Outer Mitochondrial Membrane

**DOI:** 10.1101/2025.01.20.633997

**Authors:** Sara Wong, Katherine R. Bertram, Nidhi Raghuram, Thomas Knight, Adam L. Hughes

**Affiliations:** Department of Biochemistry, University of Utah School of Medicine, Salt Lake City, UT 84112, USA

## Abstract

Lipid saturation is a key determinant of membrane function and organelle health, with changes in saturation triggering adaptive quality control mechanisms to maintain membrane integrity. Among cellular membranes, the mitochondrial outer membrane (OMM) is an important interface for many cellular functions, but how lipid saturation impacts OMM function remains unclear. Here, we show that increased intracellular unsaturated fatty acids (UFAs) remodel the OMM by promoting the formation of multilamellar mitochondrial-derived compartments (MDCs), which sequester proteins and lipids from the OMM. These effects depend on the incorporation of UFAs into membrane phospholipids, suggesting that changes in membrane bilayer composition mediate this process. Furthermore, elevated UFAs impair the assembly of the OMM protein translocase (TOM) complex, with unassembled TOM components captured into MDCs. Collectively, these findings suggest that alterations in phospholipid saturation may destabilize OMM protein complexes and trigger an adaptive response to sequester excess membrane proteins through MDC formation.

**Significance Statement:**

- Mitochondrial-derived compartments are multilamellar structures that sequester protein and lipids of the outer mitochondrial membrane in response to metabolic and membrane perturbations, but it is largely unknown how membrane fluidity influences this pathway.
- Increased levels of unsaturated phospholipids may disrupt the TOM complex, a large multi-subunit complex on the outer mitochondrial membrane, to promote the formation of mitochondrial-derived compartments, while increased levels of saturated phospholipids inhibits formation of mitochondrial-derived compartments.
- These findings reveal a link between phospholipid composition and protein stress in driving mitochondrial-derived compartment biogenesis, and thus mitochondrial quality control.

## Introduction

Lipids are building blocks of biological membranes, contributing to the structure, function, and dynamics of organelles within the cell. Cellular membranes are composed of a variety of lipid species, including phospholipids, sterols, and sphingolipids, each contributing to the unique properties of different organelles. The composition and organization of these lipids play a critical role in membrane fluidity, curvature, and protein functionality (Corin and Bowie, 2020; Klose et al., 2012; Renne and de Kroon, 2018; Sarmento et al., 2023). Among the various lipid characteristics, the degree of saturation—referring to the number of double bonds present in fatty acid chains—has important implications for membrane behavior. Saturated lipids, which lack double bonds, create more rigid and ordered membranes, whereas unsaturated lipids introduce fluidity and flexibility, allowing membranes to adapt to varying cellular demands. This balance between lipid saturation and unsaturation is essential for maintaining cellular homeostasis, especially under conditions that require membrane remodeling or stress adaptation (Ballweg and Ernst, 2017; Budin et al., 2018; Ernst et al., 2016; Romanauska and Kohler, 2023).

Changes in lipid composition and saturation levels can significantly impact the function of various organelles, which are dependent on the integrity of their lipid bilayers for maintaining proper protein folding, membrane trafficking, and overall cellular function. Membranes that become too rigid or too fluid can impair protein localization and function, triggering cellular stress responses. To adapt to these changes, cells have evolved complex quality control mechanisms to maintain organelle integrity and protein homeostasis. These mechanisms include the unfolded protein response in the endoplasmic reticulum (ER) (Halbleib et al., 2017; Shyu et al., 2019; Volmer et al., 2013) and mitochondria (Melber and Haynes, 2018), autophagy (Koh et al., 2018), lipid droplet formation (Garbarino et al., 2009; Graef, 2018; Obaseki et al., 2024; Petschnigg et al., 2009), and the extraction of altered proteins from membranes via various quality control systems (den Brave et al., 2021; Phillips et al., 2020; Sardana and Emr, 2021). These adaptive responses are critical for mitigating the negative effects of altered lipid saturation, which can disrupt cellular processes and contribute to diseases associated with membrane and protein dysfunction (Pizzuto et al., 2019).

Among the various organelles, mitochondria are highly sensitive to lipid composition changes (Joshi et al., 2023; Watson et al., 1975) due to their dual-membrane structure and central role in cellular energy production, calcium regulation, and apoptosis. While mitochondria have two dynamic membranes, their roles are quite distinct (Kuhlbrandt, 2015). The inner mitochondrial membrane (IMM) houses the oxidative phosphorylation machinery, and is critical for supporting various aspects of mitochondrial metabolism. The OMM mediates essential processes such as protein import, communication with other organelles, and the regulation of immune responses and cell death signals. While lipid saturation alterations have been shown to greatly impact the structure and function of the inner mitochondrial membrane (Budin et al., 2018; Venkatraman and Budin, 2024; Venkatraman et al., 2023), less is understood about the impacts of lipid saturation on the various functions of the outer mitochondrial membrane (OMM).

A recently emerging pathway for maintaining OMM homeostasis is the formation of mitochondrial-derived compartments (MDCs). MDCs are multi-lamellar structures (Wilson et al., 2024b) that form from the OMM in response to a variety of stresses, including metabolic perturbations (Hughes et al., 2016; Schuler et al., 2021) and protein overload in the OMM (Wilson et al., 2024a). MDCs sequester and remove proteins and lipid from the OMM during these conditions. A recent study suggested that phospholipids, particularly those involved in membrane fluidity, influence MDC formation. Specifically, loss of mitochondrial phosphatidylethanolamine (PE) triggers MDC biogenesis, whereas cardiolipin (CL) depletion impairs MDC formation (Xiao et al., 2024). However, the function of these lipids in MDC formation remains incompletely understood, as does the specific role of lipid saturation in MDC biogenesis.

In this study, we investigate the impact of lipid saturation on the OMM in *Saccharomyces cerevisiae*, specifically focusing on how phospholipid unsaturation affects TOM complex assembly and the induction of MDC formation. Our results demonstrate that elevated phospholipid unsaturation alters the OMM by stimulating MDC biogenesis, and by impairing the assembly of the TOM complex. Given the known role of MDCs in sequestering hydrophobic cargo from the OMM, we propose that MDCs may act to blunt membrane stress downstream of changes in lipid saturation.

## Results

### Changes in unsaturated fatty acid levels modulate MDC biogenesis

To investigate whether changes in lipid saturation impact the OMM and stimulate remodeling via MDCs, we examined MDC formation in cells after modulating the expression of *OLE1*, the sole fatty acid desaturase in budding yeast and homolog of Stearoyl-CoA Desaturase-1 (Stukey et al., 1989; Stukey et al., 1990). Ole1 resides on the endoplasmic reticulum (ER) membrane and desaturates C16:0 and C18:0 fatty acids before their incorporation into phospholipids or storage lipids (reviewed in (Ballweg and Ernst, 2017)). We found via whole-cell lipidomic analysis that overexpressing an extra copy of *OLE1* from a *GPD* promoter (*OLE1^OE^*) increased cellular levels of di-unsaturated phospholipids in both rich (YPAD) and defined synthetic media (SD), confirming the efficacy of this approach to boost unsaturated lipids in the cell (Figure 1 A-B and Supplementary Table 1).

**Figure 1.**
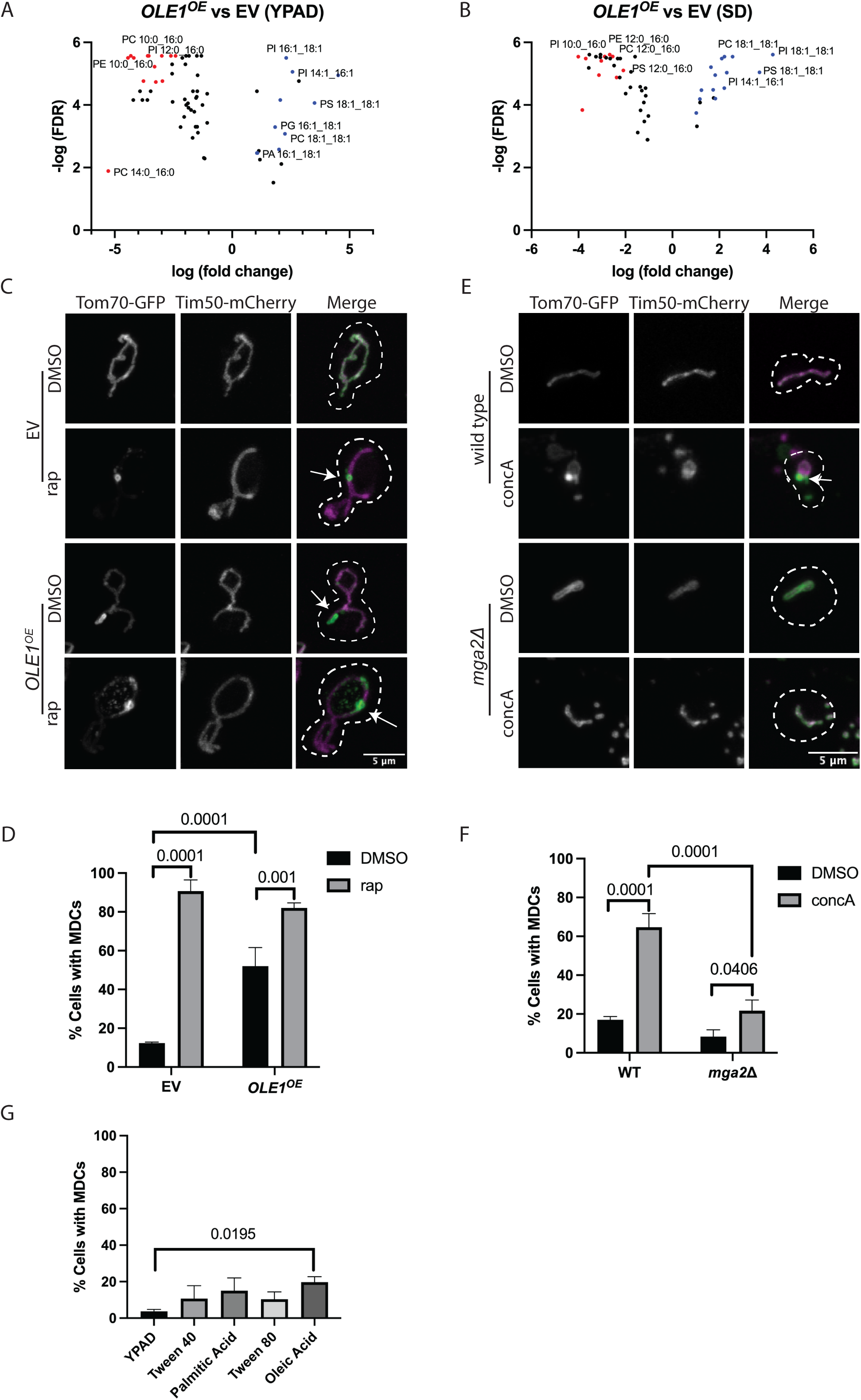
Changes in unsaturated fatty acid levels modulate MDC biogenesis. (A, B) Whole-cell lipidomic analysis of yeast overexpressing empty vector (EV) or *OLE1* (*OLE1^OE^*), grown in YPAD (A) or SD (B) media overnight. Volcano plots showing changes in lipid species. Red indicates di-saturated, blue indicates di-unsaturated, and black indicates mono-unsaturated phospholipids. (C) Super-resolution confocal fluorescence microscopy images of yeast cells overexpressing empty vector (EV) or an extra copy of *OLE1* (*OLE1^OE^*) driven by a *GPD* promoter and inserted into chromosome I, and expressing endogenously C-terminally tagged Tom70-GFP and Tim50-mCherry. Cells were treated with DMSO or rapamycin (rap) for 2 hours. Representative images of max projections. White arrows indicate MDCs. Scale Bar = 5 microns. (D) Quantification of (C) showing the percentage of cells with MDCs. n=3, 100 cells per n. Error bars = SEM and p-value as indicated by One Way ANOVA. (E) Super-resolution confocal fluorescence microscopy images of wild-type or *mga2Δ* yeast cells expressing endogenously tagged Tom70-GFP and Tim50-mCherry. Cells were treated with DMSO or rap for 2 hours. Representative images of max projections. White arrows indicate MDCs. Scale Bar = 5 microns. (F) Quantification of (E) showing the percentage of cells with MDCs. n=3, 100 cells per n. Error bars = SEM and p-value as indicated by One Way ANOVA. (G) Quantification of wild-type cells grown in YPAD containing 1% Tween 40, 1% Tween 40 + 1 mM Palmitic Acid, 1% Tween 80, or 1% Tween 80 + 1 mM Oleic Acid for 2 hours. Quantification shows the percentage of cells with MDCs. n=3, 100 cells per n. Error bars = SEM and p-value as indicated by One Way ANOVA.

We then examined whether elevated unsaturation impacts MDC levels via confocal fluorescence imaging of strains expressing well-characterized MDC markers, Tom70-GFP and Tim50-mCherry. MDCs are large subdomains derived from mitochondria that contain only OMM cargo proteins, and as such can be visualized as spherical structures that are highly enriched for certain OMM proteins including Tom70 while excluding internal mitochondrial proteins, such as Tim50 (Hughes et al., 2016). As described above, MDCs can be triggered by metabolic perturbations and protein overload stress, including elevating amino acid pools through impairment of the mTOR signaling pathway via treatment with the mTOR inhibitor rapamycin (Schuler et al., 2021). An example of rapamycin-induced MDC formation is shown in Figure 1C and quantified in Figure 1D, where MDCs are present in 90% of treated cells containing an empty-vector control. Similar to rapamycin treatment, overexpressing *OLE1* stimulated MDC formation (Figure 1C-D). 52% of *OLE1^OE^* cells formed MDCs constitutively, which was further increased with the addition of rapamycin (Figure 1C-D). Likewise, treatment with another well-characterized metabolic MDC inducer, concanamycin A (concA), a potent inhibitor of vacuole acidification that triggers MDCs through amino acid perturbation (Hughes et al., 2016; Schuler et al., 2021), also further elevated MDC levels in *OLE1^OE^*cells, indicating additive effects across these perturbations (Supplementary Figure 1A). Importantly, these Tom70 structures formed by *OLE1^OE^*required the mitochondrial-localized GTPase Gem1 for formation, which was previously shown to be required for MDC biogenesis (English et al., 2020) (Supplementary Figure 1B). Thus, these structures in *OLE1^OE^* cells are indeed MDCs, based on their characteristics and genetic requirements.

As an orthogonal approach, we tested whether expression of an activated form of *SPT23*, a transcription factor that regulates expression of *OLE1*, could also stimulate MDC formation (Zhang et al., 1999). Spt23 resides on the ER membrane and senses membrane fluidity. When membranes are rigid, Spt23 is cleaved from the membrane, translocates to the nucleus, and stimulates expression of *OLE1* (reviewed in (Ballweg and Ernst, 2017)). Expression of an active, truncated form of Spt23 (Belgareh-Touze et al., 2017) also induced constitutive formation of MDCs, similar to overexpression of *OLE1* (Supplementary Figure 1C). These data suggest that increasing *OLE1* expression, either via a constitutive promoter or by activating a transcription factor that controls its expression, induces MDC formation.

We next tested if reducing *OLE1* expression impacts MDC formation. Because *OLE1* is an essential gene and not easily depleted, we deleted *MGA2*, another transcription factor that regulates *OLE1* expression (Zhang et al., 1999) in order to reduce Ole1 levels in the cell. In contrast to *OLE1* overexpression, deletion of *MGA2* did not stimulate MDC formation (Figure 1E-F). In fact, loss of Mga2 impaired MDC formation. While 65% of wild type cells formed MDCs when treated with concA, only 22% of *mga2Δ* mutant cells formed MDCs under this condition (Figure 1E-F). Overexpression of *OLE1* from the constitutive *GPD* promoter in *mga2Δ* mutant cells restored MDC formation (Supplementary Figure 1D), confirming that MDC inhibition results from reduced expression of *OLE1* and not other transcriptional targets of Mga2. Finally, we found that acute addition of fatty acids to the growth medium did not strongly activate MDC formation (Figure 1G). This lack of response may stem from the difficulty of delivering fatty acids to yeast, and because excess fatty acids, especially dietary fatty acids, are often shunted to lipid droplets for storage (reviewed in (Zadoorian et al., 2023)). Overall, our data suggest that UFA synthesis regulates MDC formation— elevated lipid unsaturation activates MDC formation, whereas increased saturation suppresses MDC biogenesis.

### The acyltransferase Sct1 antagonizes the effect of Ole1 on MDC biogenesis

Previous studies found that the acyltransferase, Sct1, competes with Ole1 for substrates, and that Sct1 preferentially incorporates saturated acyl chains into phospholipids (De Smet et al., 2012). Thus, overexpression of *SCT1* leads to more saturated phospholipids in cells and co-overexpression of *SCT1* and *OLE1* largely restores lipid balance (De Smet et al., 2012), with a modest shift towards elevated UFAs (Supplementary Figure S2A-B and Supplementary Table 2). Based on our data that MDC formation is suppressed in *mga2Δ* cells due to reduced *OLE1* expression (Figure 1E-F), we hypothesized that overexpression of *SCT1* may suppress MDCs, and that co-overexpression of *SCT1* and *OLE1* would restore MDC formation to normal levels. Indeed, we found that overexpression of *SCT1* (*SCT1^OE^*) modestly reduced MDCs triggered by concA or *OLE1* overexpression (Figure 2A-B). MDC suppression by *SCT1* overexpression was not as apparent in rapamycin treated cells, based on the percentage of cells that formed MDCs (Figure 2B). However, we noted that *OLE1*^OE^ increased the average diameter of MDCs in rapamycin treated cells, and that *SCT1^OE^*prevented this increase, suggesting Sct1 also has suppressive effects in the presence of rapamycin (Figure 2C-D). Finally, we found that deletion of *SCT1*, which causes an increase in unsaturated phospholipids (De Smet et al., 2012), triggered constitutive MDC formation comparable to *OLE1^OE^* (Figure 2E). Thus, Sct1 and Ole1 appear to antagonize one another in the regulation of MDC formation, and increasing the level of saturated fatty acid chains incorporated into phospholipids suppresses MDC biogenesis.

**Figure 2.**
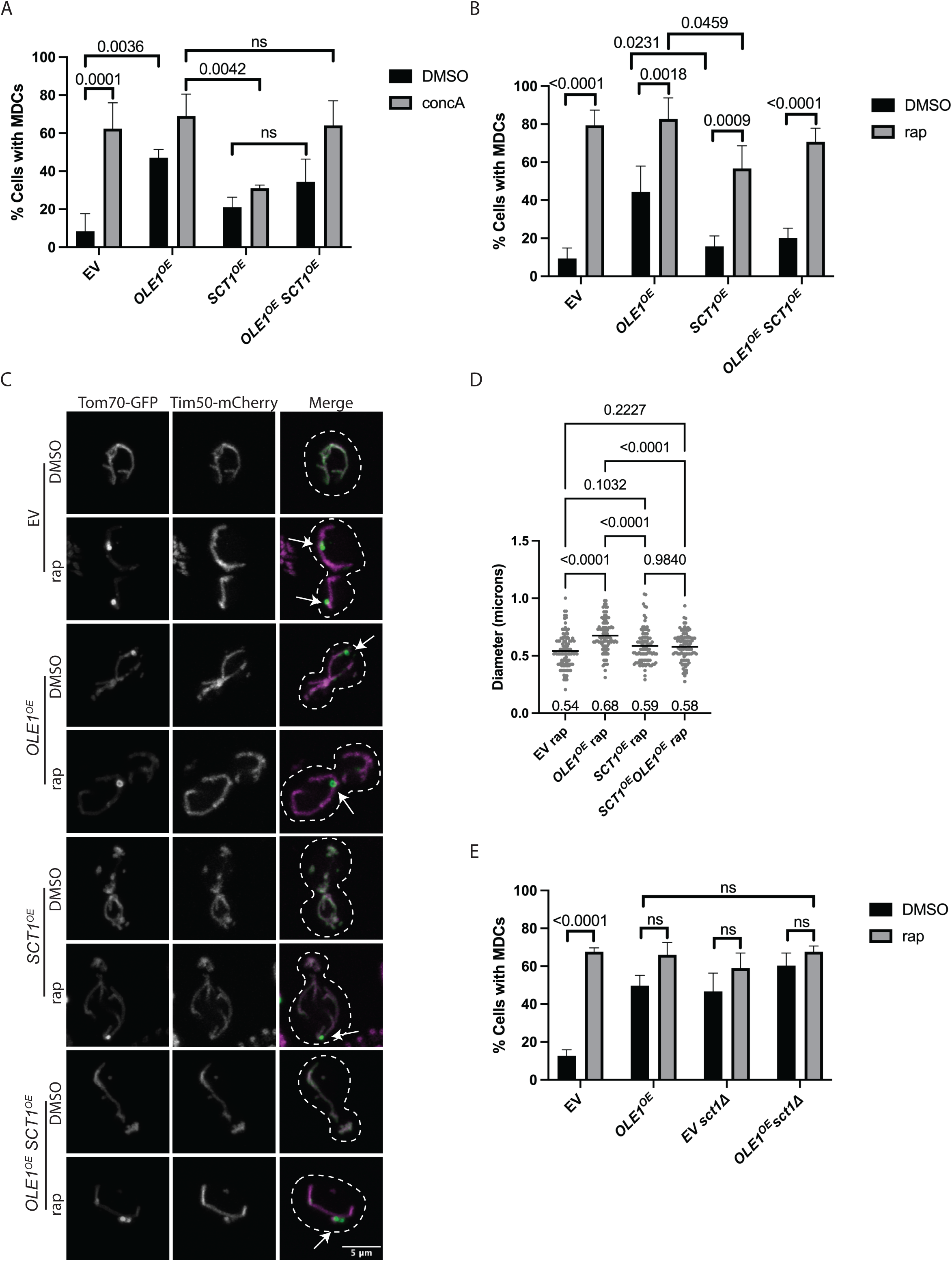
The acyltransferase Sct1 antagonizes the effect of Ole1 on MDC biogenesis. (A, B) Quantification of the percentage of yeast cells overexpressing empty vector (EV), *OLE1* (*OLE1^OE^*), *SCT1* (*SCT1^OE^*), or both (*OLE1^OE^SCT1^OE^*) exhibiting MDCs. Cells were treated with DMSO, concA (A) or rap (B) for 2 hours. n=3, 100 cells per n. Error bars = SEM and p-value as indicated by One Way ANOVA. (C) Super-resolution confocal fluorescence microscopy images of yeast cells overexpressing empty vector (EV), *OLE1* (*OLE1^OE^*), *SCT1* (*SCT1^OE^*), or both (*OLE1^OE^SCT1^OE^*) from a *GPD* promoter in Chromosome I, and endogenously tagged Tom70-GFP and Tim50-mCherry. Cells were treated with DMSO or rap for 2 hours. Representative images of max projections. White arrows indicate MDCs. Scale Bar = 5 microns. (D) Quantification of (C) showing MDC diameter with mean diameter indicated along x-axis. n=3, 30-35 MDCs per n for a total of 100 MDCs. p-value as indicated by One Way ANOVA. (E) Quantification of yeast cells overexpressing empty vector (EV) or *OLE1* (*OLE1^OE^*) in wild-type or *sct1Δ* cells. Cells were treated with DMSO or rap for 2 hours. Quantification shows the percentage of cells with MDCs. n=3, 100 cells per n. Error bars = SEM and p-value as indicated by One Way ANOVA.

### Phospholipid synthesis is required for UFAs to induce MDC formation

UFAs can be incorporated into phospholipids as part of biological membranes, or alternatively stored in sterol esters or triglycerides (TGs) in lipid droplets. We next tested whether shunting UFAs preferentially into phospholipids or TGs affects their ability to stimulate MDC formation. To do this, we altered genes in the conserved Pah1/LIPIN pathway to shunt UFAs preferentially into either TG or phospholipids. Pah1 is the yeast phosphatidate phosphatase, which catalyzes the conversion of phosphatidic acid (PA) to diacylglycerol (DG) (Adeyo et al., 2011; Han et al., 2006; Irie et al., 1993; Peterfy et al., 2001). Because PA is the precursor for phospholipids, Pah1 activity promotes FA incorporation into storage lipids, limiting phospholipid biosynthesis downstream of PA. Pah1 can be dephosphorylated and activated by the Nem1-Spo7 complex to promote synthesis of DG from PA, and thus lipid droplet biogenesis (O’Hara et al., 2006; Santos-Rosa et al., 2005). In contrast, Ice2 is a negative regulator of Pah1 that prevents this dephosphorylation and thus promotes the synthesis PA and downstream phospholipids (Papagiannidis et al., 2021). Thus, *ice2Δ* cells have high Pah1 activity and elevated levels of storage lipids, while *nem1Δ* cells exhibit increased phospholipids and lower storage lipids.

We examined MDC formation in *OLE1^OE^* cells lacking *ICE2*, and found that loss of *ICE2*, and thus preferential incorporation of UFAs into DG and TG for storage, suppressed MDC formation (Figure 3A-B). To test whether loss of the positive regulator of Pah1, Nem1, could enhance MDC formation, we conducted experiments in synthetic medium (SD), which we previously showed lowers MDC formation in cells through incompletely understood mechanisms likely linked to intracellular amino acid load and changes in lipid composition (Schuler et al., 2021). Indeed, *OLE1^OE^*-induced MDCs were blunted in SD medium (Figure 3C), even though *OLE1* overexpression in SD medium still increased UFA levels (Figure 1B). Interestingly, deleting *NEM1* in *OLE1^OE^* cells and shunting more UFAs into phospholipids significantly enhanced MDC formation in SD medium (Figure 3D-E). Altogether, these results suggest that MDCs are sensitive to the levels of UFAs in membrane phospholipids, and that shunting UFAs into storage lipids prevents them from activating the MDC pathway.

**Figure 3.**
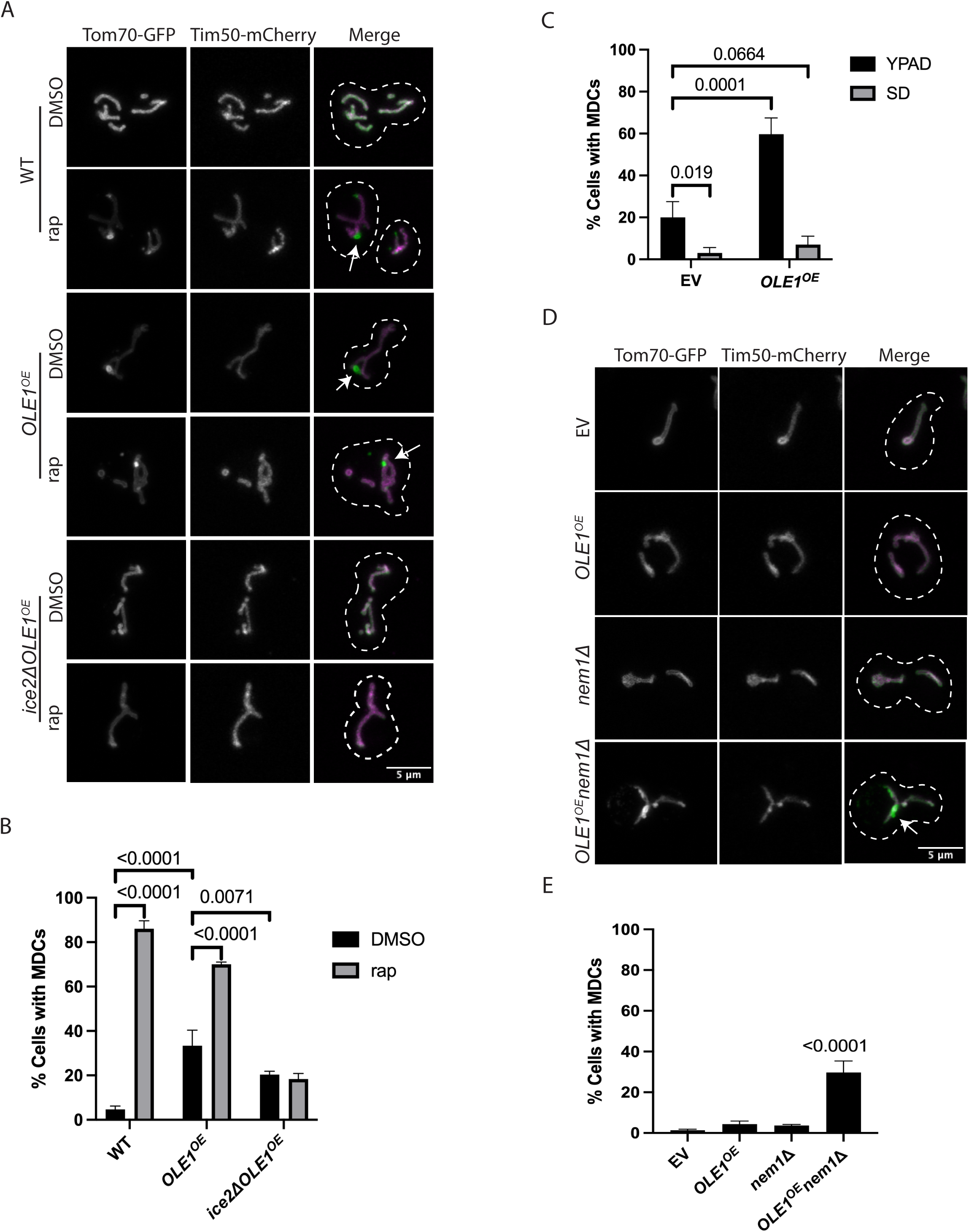
Phospholipid synthesis is required for UFAs to induce MDC formation. (A) Super-resolution confocal fluorescence microscopy images of wild-type (WT), *OLE1^OE^*, or *ice2Δ OLE1^OE^* cells expressing endogenously tagged Tom70-GFP and Tim50-mCherry. Cells were treated with DMSO or rap for 2 hours. Representative images of max projections. White arrows indicate MDCs. Scale Bar = 5 microns. (B) Quantification of (A) showing the percentage of cells with MDCs. n=3, 100 cells per n. Error bars = SEM and p-value as indicated by One Way ANOVA. (C) Quantification of the percentage of cells with MDCs for yeast overexpressing EV or *OLE1* and grown in YPAD or SD media. n=3, 100 cells per n. Error bars = SEM and p-value as indicated by One Way ANOVA. (D) Super-resolution confocal fluorescence microscopy images of yeast cells overexpressing empty vector (EV) or *OLE1* (*OLE1^OE^*) in wild-type or *nem1Δ* cells with endogenously tagged Tom70-GFP and Tim50-mCherry. Representative images of max projections. White arrows indicate MDCs. Scale Bar = 5 microns. (E) Quantification of (D) showing the percentage of cells with MDCs. n=3, 100 cells per n. Error bars = SEM and p-value as indicated by One Way ANOVA.

### Increased phospholipid unsaturation may cause protein stress on the outer mitochondrial membrane

Finally, we wanted to understand how elevated UFAs in membrane phospholipids may trigger MDC formation. To date, MDCs have been shown to be stimulated by changes in intracellular metabolites, including coupled alterations in intracellular amino acids and mitochondrial TCA cycle metabolites, as well as alterations in phospholipid species and protein overload or protein mistargeting stress in the OMM (Raghuram and Hughes, 2024; Schuler et al., 2021; Wilson et al., 2024a; Xiao et al., 2024). Whole-cell metabolite analysis in cells overexpressing *OLE1* showed that TCA cycle metabolites were not changed in *OLE1^OE^* cells compared to an empty vector control (Supplementary Figure 3 and Supplementary Table 3). Thus, it does not appear that UFAs stimulate MDCs via altering the TCA cycle, which is the current model as to how elevated amino acids (via rapamycin and concA treatment) stimulate the pathway (Raghuram and Hughes, 2024).

Based on this result, we instead explored whether any links exist between UFAs and protein stress in the OMM—another robust MDC inducer. It was recently found that MDC formation may be induced by mistargeted and excess proteins on the OMM, and that loss of Msp1—a AAA-ATPase on the mitochondrial outer membrane that helps to remove mistargeted tail-anchored (TA) proteins in yeast (Chen et al., 2014; Matsumoto et al., 2019; Schuldiner et al., 2008), arrested precursor proteins in *C. elegans* (Basch et al., 2020), and TOM complexes during import stress in both yeast (Weidberg and Amon, 2018) and mammals (Kim et al., 2024)—enhanced MDC formation (Wilson et al., 2024a). We found that deletion of *MSP1* resulted in 40% of cells forming MDCs constitutively, consistent with previous reports (Figure 4A, 4B) (Wilson et al., 2024a). Additionally, *MSP1* deletion increased the penetrance of MDCs in *OLE1^OE^* cells, and enhanced the average diameter of MDCs in *OLE1^OE^*cells treated with rapamycin (Figure 4A-C). These results suggested a potential interplay between *OLE1^OE^* and protein overload stress in the OMM.

**Figure 4.**
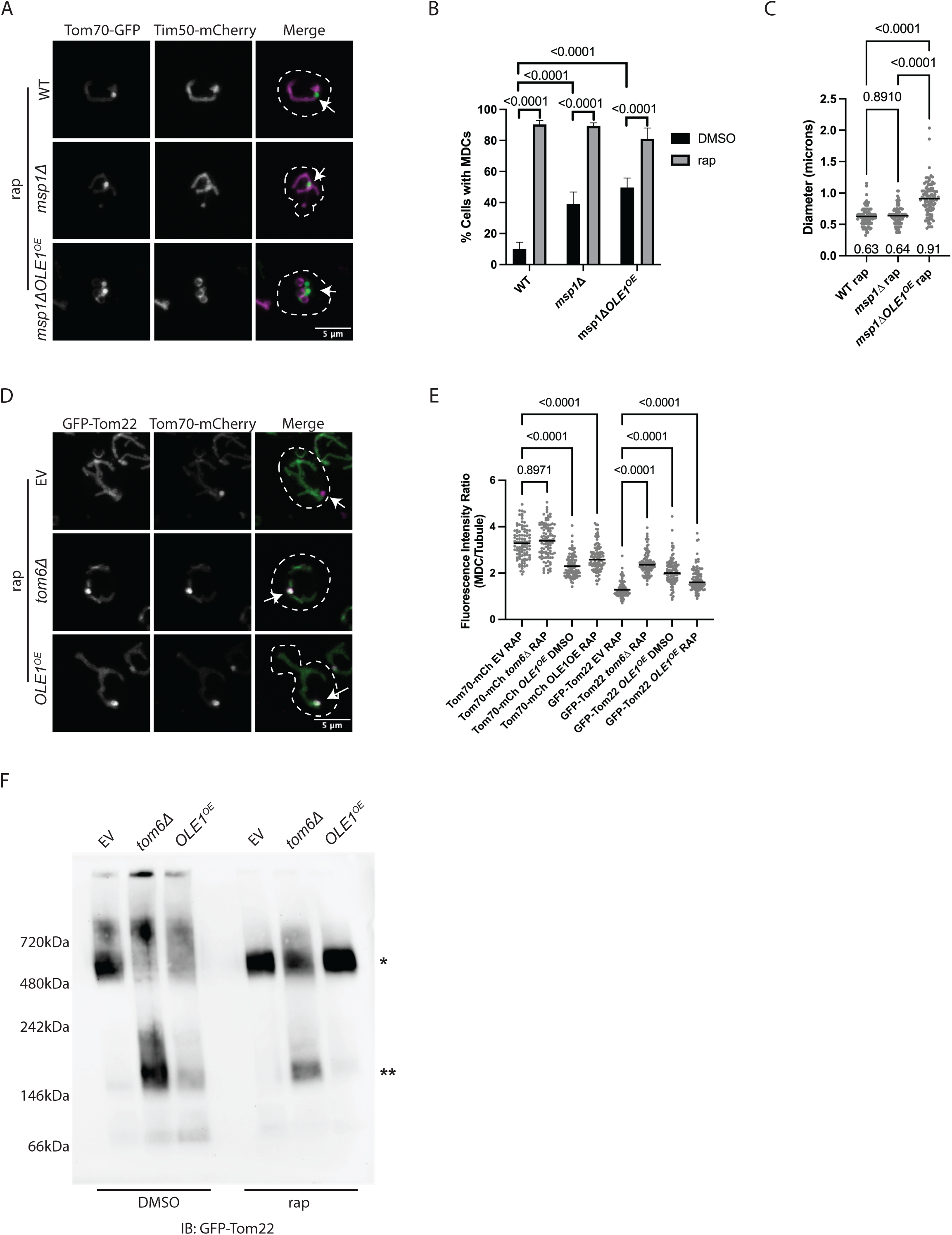
Increased phospholipid unsaturation perturbs protein complexes on the outer mitochondrial membrane. (A) Super-resolution confocal fluorescence microscopy images of wild-type (WT), *msp1Δ*, or *msp1ΔOLE1^OE^*cells expressing endogenously tagged Tom70-GFP and Tim50-mCherry, and treated with rap for 2 hours. Representative images of max projections. White arrows indicate MDCs. Scale Bar = 5 microns. (B) Quantification of (A) showing the percentage of cells with MDCs. n=3, 100 cells per n. Error bars = SEM and p-value as indicated by One Way ANOVA. (C) Quantification of (A) showing MDC diameter, with mean diameter below scatterplot. n=3, 30-35 MDCs per n for a total of 100 MDCs. p-value as indicated by One Way ANOVA. (D) Super-resolution confocal fluorescence microscopy images of EV, *tom6Δ*, or *OLE1^OE^*cells expressing endogenously tagged Tom70-mCherry and GFP-Tom22, and treated with rap for 2 hours. Representative images of max projections. White arrows indicate MDCs. Scale Bar = 5 microns. (E) Quantification of (D) showing fluorescence intensity of the MDC compared to the mitochondrial tubule. n=3, 100 MDCs per n. Bar shows mean. p-value as indicated by One Way ANOVA. (F) Blue-Native PAGE analysis of mitochondria isolated from EV, *tom6Δ*, or *OLE1^OE^*mutant cells expressing endogenously tagged Tom70-mCherry and GFP-Tom22, and treated with DMSO or rap for 2 hours. Immunoblot for GFP. **\*** indicates intact TOM complex, **\*\*** indicates disassembled TOM complex. Representative of n=2.

To investigate this link further, we first tested whether elevated unsaturated phospholipids caused mistargeting of ER-localized TA proteins to the OMM. TA mistargeting in this manner occurs in cells with a perturbed GET pathway, which targets TA proteins to the ER (Schuldiner et al., 2008). Importantly, it was previously shown that loss of GET pathway components *GET1* and *GET2* increased MDC formation and led to capture of mistargeting TA proteins into MDCs (Wilson et al., 2024a). However, unlike loss of the GET pathway, we found no evidence of TA mistargeting in *OLE1^OE^* cells, as the model TA substrate GFP-Ubc6 did not localize to mitochondria or MDCs in *OLE1^OE^*cells (Supplementary Figure 4A). These results suggest that elevated membrane unsaturation does not trigger MDCs via TA protein overload stress in the OMM.

Next, we considered the possibility that elevated lipid unsaturation may impact the TOM import machinery on the OMM, which could lead to protein overload in the OMM. Previous studies found that most subunits of the TOM complex are excluded from MDCs in rapamycin or concA treated cells, unless the assembly of the complex is disrupted, such as in cells lacking the small TOM subunit, Tom6 (Dekker et al., 1998; Wilson et al., 2024a). Indeed, as previously reported we found that the TOM complex subunit GFP-Tom22 was excluded from rapamycin-induced MDCs, but incorporated into MDCs in cells lacking *TOM6* (Figure 4D-E). Importantly, we found that GFP-Tom22 was enriched in MDCs in *OLE1^OE^* cells, similar to a *tom6Δ* mutant (Figure 4D-E). Other TOM complex components that are normally excluded from MDCs in rapamycin treatment, including GFP-Tom5 and GFP-Tom7, were also enriched in MDCs in *OLE1^OE^*cells, similar to a *tom6Δ* mutant (Supplementary Figure 4B-C). These results suggest that increased phospholipid unsaturation may disrupt the assembly of the TOM complex in the OMM, leading to sequestration of unassembled TOM components into MDCs.

To examine this possibility further, we carried out Blue-Native PAGE analysis to examine TOM complex assembly upon overexpression of *OLE1*. By immunblotting for TOM complex subunit GFP-Tom22, we found that levels of intact TOM complex were reduced in cells overexpressing *OLE1^OE^* with an increase in free Tom22, similar to what occurs in a *tom6Δ* mutant (Figure 4F). For currently unknown reasons, this effect is partially blunted upon rapamycin treatment, possibly due to a decrease in *OLE1* expression during rapamycin treatment (Supplementary Figure 4D) (Iesmantavicius et al., 2014). In contrast to *OLE1^OE^*, overexpression of *SCT1* did not lead to an enrichment of the TOM complex subunit GFP-Tom7 in MDCs (Supplementary Figure 4E), and did not cause altered assembly of the TOM complex (Supplementary Figure 4F), suggesting that these effects are specific to elevated membrane unsaturation. Collectively, these results suggest that elevated membrane phospholipid unsaturation alters the assembly of the TOM complex, causing an increase in unassembled TOM complex components in the OMM and sequestration of these components into MDCs.

## Discussion

Lipid saturation is tightly regulated, and defects in maintaining proper saturation levels have been shown to have a multitude of effects on cells. While there are many positive and protective impacts of increased lipid unsaturation (Akazawa et al., 2010; Dalla Valle et al., 2019; Fang et al., 2017; Huang et al., 2018; Li et al., 2019; Miller et al., 2005; Nasution et al., 2017; Tuthill Ii et al., 2021), upregulated unsaturated lipids can also lead to lipotoxicity and are correlated with several types of diseases (Abd Alla et al., 2021; AM et al., 2017; Balatskyi and Dobrzyn, 2023; Kikuchi and Tsukamoto, 2020; Kim et al., 2011; Liu et al., 2011; Paton and Ntambi, 2009; Yamamoto and Sano, 2022).

In this study, we sought to better understand how elevated levels of UFAs in cells specifically impact organelle homeostasis, with a focus on mitochondria. Our results identified a previously unknown role for UFAs in modifying the OMM, triggering dissociation of TOM complex subunits and stimulating biogenesis of OMM-derived multilamellar compartments, or MDCs. Interestingly, UFA-induced MDC formation was suppressed by shifting the distribution of UFAs into storage lipids and away from phospholipids, suggesting that the stimulatory impact of UFAs on MDCs occurs through their effect on phospholipid-containing membrane bilayers. In contrast to unsaturated lipids, an increase in saturation did not stimulate MDCs, suggesting that OMM-remodeling via MDCs is not a general response to changes in membrane fluidity.

Our data that UFAs perturb the assembly of the TOM complex in the OMM suggests that unsaturated lipids may induce protein stress at the OMM, potentially disrupting protein-protein interactions required for TOM complex stability. Because prior studies showed that MDCs can be triggered by excess hydrophobic cargo in the OMM (Wilson et al., 2024a), we propose that MDCs stimulated by UFAs may act as an adaptive response to sequester excess or mis-localized proteins in the OMM generated by alterations in the lipid bilayer. Another more speculative possibility is that MDCs may act to sequester specific lipid species from the OMM, thus helping to maintain membrane integrity. While we find that disrupted TOM complex components are sequestered into UFA-induced MDCs, the extent to which other OMM protein complexes are affected by elevated UFAs and whether they become targeted to MDCs remains unclear. Future studies investigating whether specific lipids can be incorporated into MDCs, and the breadth of proteins affected by lipid unsaturation stress – including protein alterations and membrane remodeling responses at other organelles - will be important for understanding the full scope of UFA-induced cellular stress and the function of MDCs in cells.

In conclusion, our findings uncover a new mechanism by which elevated lipid unsaturation induces stress at the OMM, and suggest that MDCs may play an important role in adapting to lipid-induced membrane stress at the mitochondria. In addition to new insights into the impacts of elevated UFAs on cellular homeostasis, this work also adds to our growing understanding of the role of MDCs as a mitochondrial adaptation pathway. It now appears that MDCs sequester portions of the OMM in response to a variety of stressors—including metabolic perturbations, changes in lipid composition, and alterations in OMM protein load and/or composition. Whether these stimulatory routes are mechanistically connected and how forming an MDC ultimately modifies mitochondrial health under these conditions remains unclear and are important areas for future investigation.

## DATA AVAILABILITY

All reagents used in this study are available upon request. All other data reported in this paper will be shared by the lead contact upon request. This paper does not report original code. Any additional information required to reanalyze the data reported in this paper is available from the lead contact upon request.

## ACKNOWLEDGEMENTS

We thank members of the Adam L. Hughes lab for their insightful discussions. We thank Zachary Wilson, PhD for construction of the Ole1-GFP yeast strain. We thank Kylie N. Jacobs, PhD for construction of the *ICE2* and *NEM1* mutants. Metabolomics and lipidomics analysis was performed at the Metabolomics Core Facility at the University of Utah. We thank Dr. John Alan Maschek for his help with lipidomics analysis. We thank Dan Cuthbertson of Agilent Technologies for assistance in implementing iterative exclusion in the tandem mass spectrometry experiments. S. Wong was supported by a National Institutes of Health Training Grant HL007576. Research was also supported by National Institutes of Health grants GM119694 and AG061376 (A.L.H).

## AUTHOR CONTRIBUTIONS

Conceptualization, S. Wong and A.L. Hughes; methodology, S. Wong and N. Raghuram; formal analysis, S. Wong, K.R. Bertram, and N. Raghuram; investigation, S. Wong, K.R. Bertram, N. Raghuram, and T. Knight; writing – original draft, S. Wong; writing – review and editing, S. Wong and A.L. Hughes; visualization, S. Wong and K. Bertram; supervision, A.L. Hughes; funding acquisition, S. Wong and A.L. Hughes.

## CONTACT FOR REAGENT AND RESOURCE SHARING

Further information and requests for resources and reagents should be directed to and will be fulfilled by the Lead Contact, Adam L. Hughes. All unique/stable reagents generated in this study are available from the Lead Contact without restrictions.

---

**Supplementary Figure 1.**
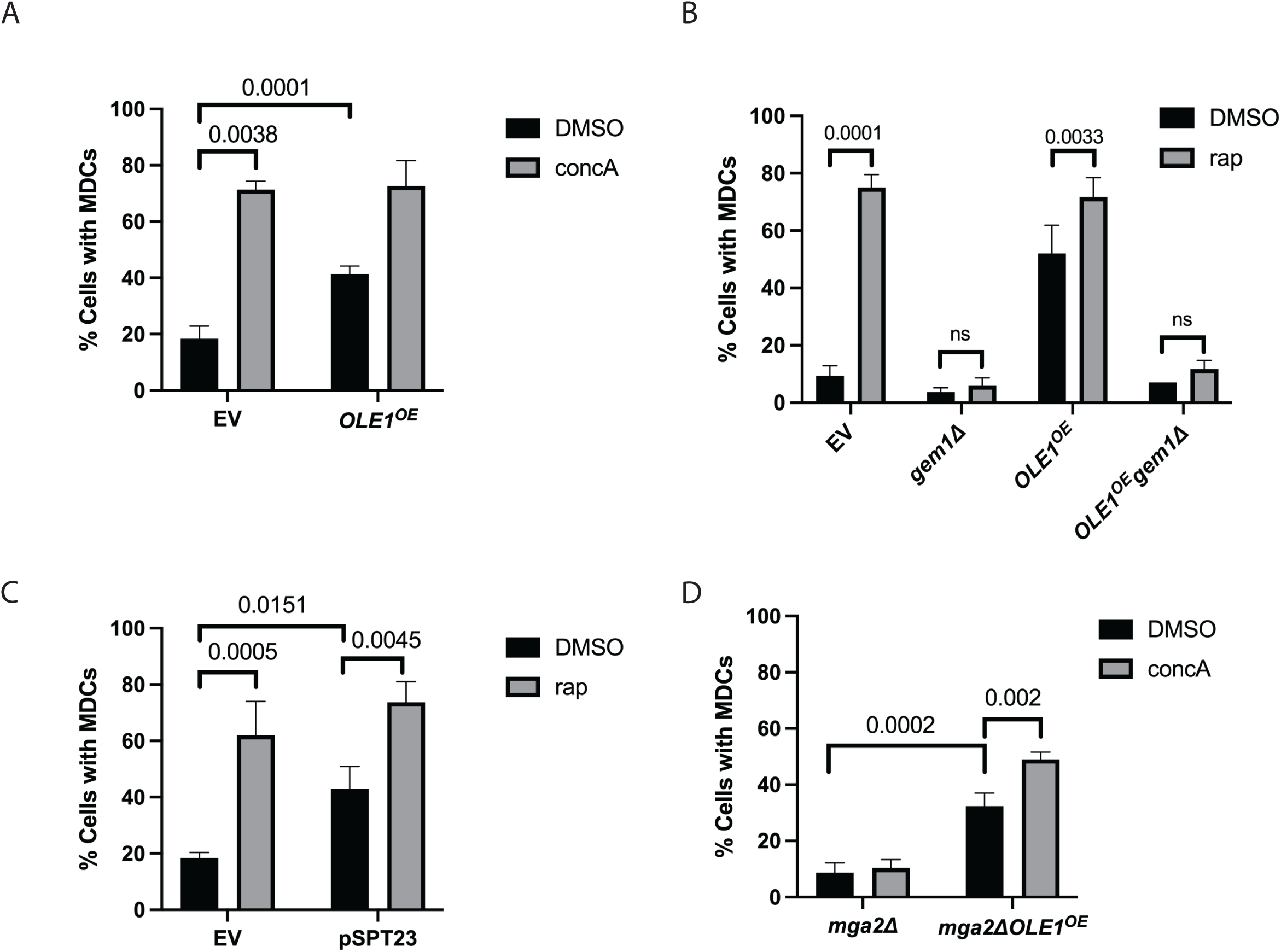
Changes in unsaturated fatty acid levels modulate MDC biogenesis, related to Figure 1. (A) Quantification of the percentage of cells with MDCs in yeast overexpressing empty vector (EV) or *OLE1* (*OLE1^OE^*) and treated with DMSO or concanamycin A (concA) for 2 hours. n=3, 100 cells per n. Error bars = SEM and p-value as indicated by One Way ANOVA. (B) Quantification of the percentage of cells with MDCs in the indicated strains treated with DMSO or rap for 2 hours. n=3, 100 cells per n. Error bars = SEM and p-value as indicated by One Way ANOVA. (C) Quantification of MDCs numbers in yeast cells expressing empty vector (EV) or Spt23(1-686) (pSTP23) from a plasmid and treated with DMSO or rap for 2 hours. n=3, 100 cells per n. Error bars = SEM and p-value as indicated by One Way ANOVA. (D) Quantification of MDC numbers in *mga2Δ* cells overexpressing empty vector (*mga2Δ*) or *OLE1* (*mga2ΔOLE1^OE^*) and treated with DMSO or concA for 2 hours. n=3, 100 cells per n. Error bars = SEM and p-value as indicated by One Way ANOVA.

**Supplementary Figure 2.**
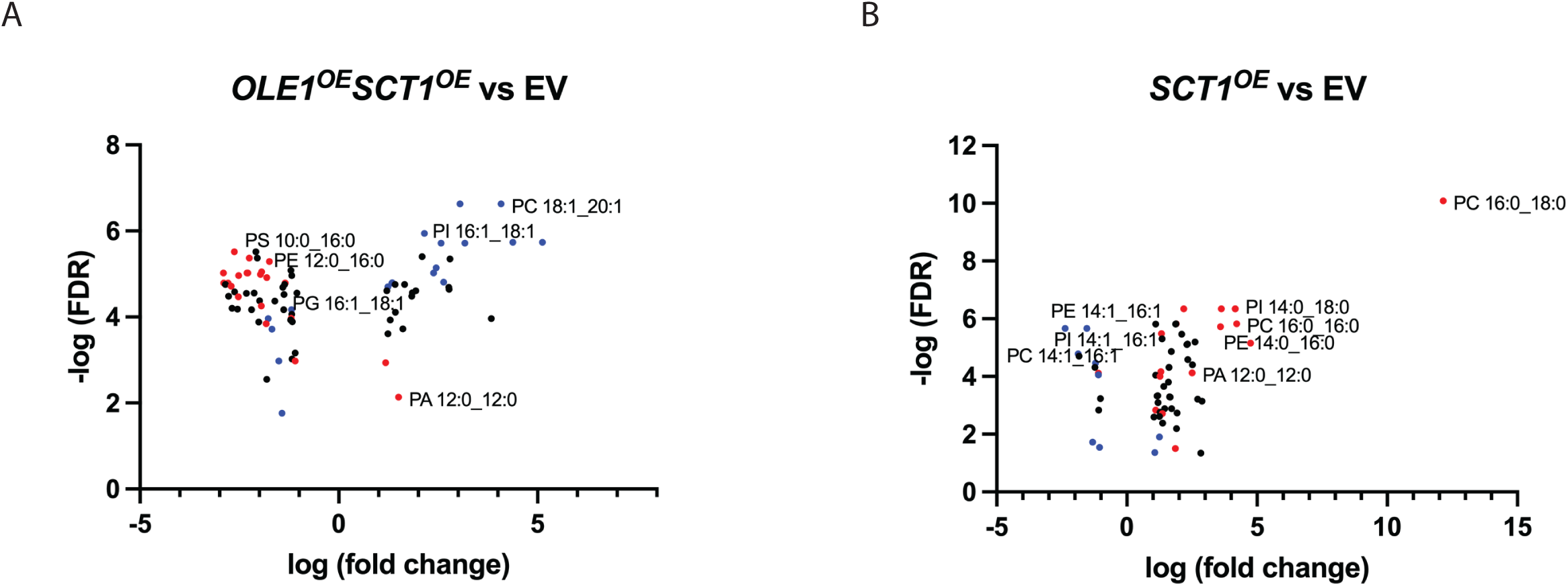
Phospholipid saturation profiles are changed in strains with altered expression of Ole1 and Sct1, related to Figure 2. Whole-cell lipidomic analysis of yeast cells overexpressing empty vector (EV), *OLE1* (*OLE1^OE^*), *SCT1* (*SCT1^OE^*), or both (*OLE1^OE^ SCT1^OE^*). Volcano plots showing changes in lipid species. Red indicates di-saturated, blue indicates di-unsaturated, and black indicates mono-unsaturated phospholipids. (A) Compares *OLE1^OE^ SCT1^OE^* and EV, (B) compares *SCT1^OE^* and EV.

**Supplementary Figure 3.**
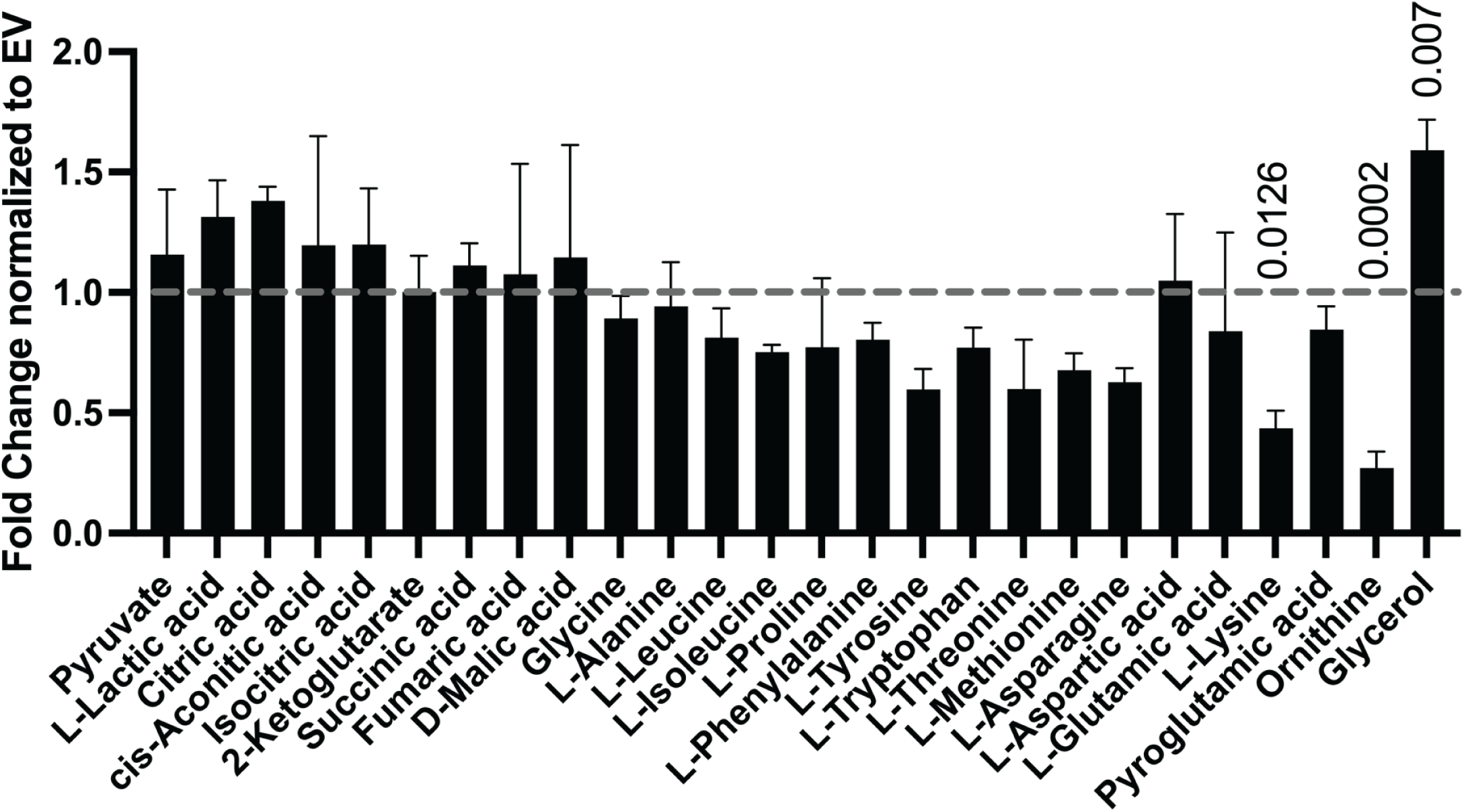
Overexpression of *OLE1* does not alter the abundance of TCA cycle metabolites. Whole cell steady-state metabolomics of *OLE1^OE^* cells, normalized to EV (dashed line). n=4. Error bars = SEM. p-value as indicated by t-test.

**Supplementary Figure 4.**
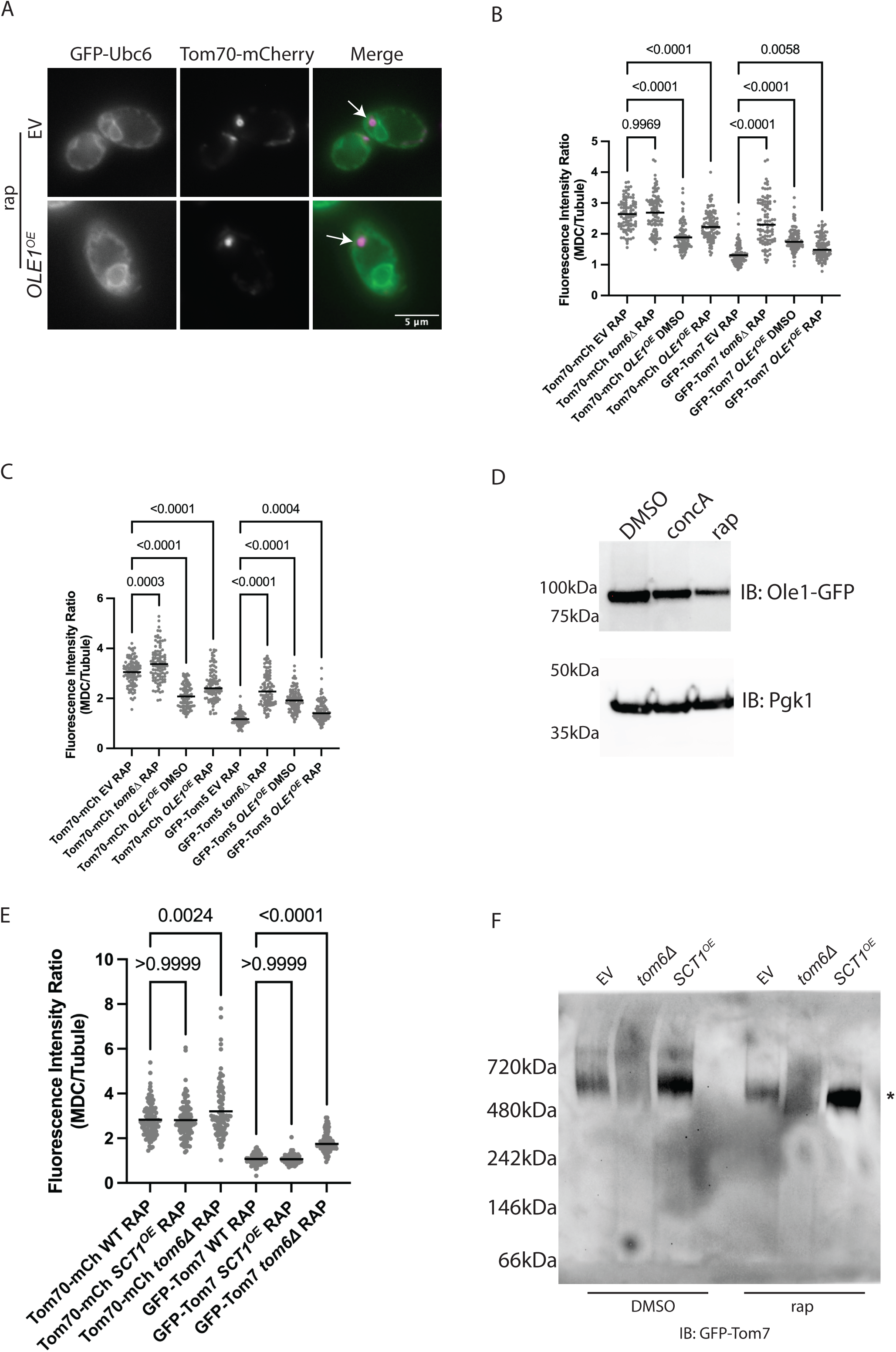
Increased phospholipid unsaturation perturbs protein complexes on the outer mitochondrial membrane, related to Figure 4. (A) Super-resolution confocal fluorescence microscopy images of EV or *OLE1^OE^*cells expressing endogenously tagged Tom70-mCherry and GFP-Ubc6, and treated with rap for 2 hours. Representative images of max projections. White arrows indicate MDCs. Scale Bar = 5 microns. (B) Quantification of EV, *tom6Δ*, or *SCT1^OE^*cells expressing endogenously tagged Tom70-mCherry and GFP-Tom7, and treated with rap for 2 hours. Quantification shows fluorescence intensity of the MDC to mitochondrial tubule. n=3, 100 MDCs per n. Bar shows mean. p-value as indicated by One Way ANOVA. (C) Quantification of EV, *tom6Δ*, or *SCT1^OE^* cells expressing endogenously tagged Tom70-mCherry and GFP-Tom5, and treated with rap for 2 hours. Quantification shows fluorescence intensity of the MDC to mitochondrial tubule. n=3, 100 MDCs per n. Bar shows mean. p-value as indicated by One Way ANOVA. (D) Whole cell lysates of yeast endogenously expressing Ole1-GFP and treated with DMSO, concA, or rap for 2 hours. Lysates were analyzed by western blot and immunoblotted for GFP and loading control Pgk1. Representative of n=3. (E) Quantification of EV, *tom6Δ*, or *SCT1^OE^*cells expressing endogenously tagged Tom70-mCherry and GFP-Tom7, and treated with rap for 2 hours. Quantification shows fluorescence intensity of the MDC to mitochondrial tubule. n=3, 100 MDCs per n. Bar indicates mean. (F) Blue-Native PAGE analysis of mitochondria isolated from EV, *tom6Δ*, or *SCT1^OE^*cells expressing endogenously tagged Tom70-mCherry and GFP-Tom7, and treated with DMSO or rap for 2 hours. Immunoblot for GFP. * indicates intact TOM complex. Representative of n=2.

## Supplementary Tables

Supplementary Table 1 contains the MetaboAnalyst Input data for Lipidomics experiments of EV and *OLE1^OE^* cells related to Figure 1A-B. Supplementary Table 2 contains the MetaboAnalyst Input data for Lipidomics experiments of EV and *SCT1^OE^ OLE1^OE^* cells related to Supplementary Figure 2. Supplementary Table 3 contains the MetaboAnalyst Input data for Metabolomics experiments of EV and *OLE1^OE^*cells related to Supplementary Figure 3. Supplementary Table 4 lists the yeast strains used in this study. Supplementary Table 5 lists the plasmids used in this study. Supplementary Table 6 lists the oligonucleotides used in this study.

## METHODS

### Yeast Strains, Plasmids, and Reagents

All yeast strains are derivatives of *Saccharomyces cerevisiae* S288C (BY) and listed in Supplementary Table 4. Deletion strains were created by PCR mediated gene replacement using the pRS series of vectors, as previously described (Brachmann et al., 1998). Endogenously tagged fluorescent proteins were created by PCR mediated epitope tagging, as previously described (Longtine et al., 1998)(Sheff and Thorn, 2004). Plasmids for *GPD* driven expression of *OLE1 and SCT1* were generated by Gateway mediated transfer of corresponding ORF (Harvard Institute of Proteomics) from pDONR 201/221 into a pAG306-ccdB chromosome I (Hughes and Gottschling, 2012) using Gateway LR Clonase II enzyme mix (ThermoFisher) according to the manufacturer’s instructions. To integrate the resulting expression plasmid into yeast chromosome I (199456-199457), pAG306GPD-ORF chromosome I was digested with NotI. All insert sequences were verified by Azenta/Genewiz sequencing. Plasmids and reagents used in this study are listed in Supplementary Table 5. The pSPT23 plasmid was a gift from Mickael M. Cohen. Correct integrations were confirmed by a combination of colony PCR across the chromosomal integration site, correctly localized expression of fluorophore by microscopy, and/or presence of an epitope tag of the correct size by western blot. Oligos used in strain and plasmid construction are listed in Supplementary Table 6.

### Yeast Culture

Yeast were grown exponentially for 15-16 hours at 30°C to an OD600 of 0.2-1. Cells were cultured in YPAD medium (1% yeast extract, 2% peptone, 0.005% adenine, 2% glucose) or synthetic defined (SD) media (0.67% nitrogen base without amino acids, 2% glucose, 0.072 g/L each adenine, alanine, arginine, asparagine, aspartic acid, cysteine, glutamic acid, glutamine, glycine, histidine, myo-inositol, isoleucine, lysine, methionine, phenylalanine, proline, serine, threonine, tryptophan, tyrosine, uracil, valine, 0.369g/L leucine, and 0.007 g/L para-aminobenzoic acid). If indicated, cells were treated with 200 nM rapamycin or 500 nM concanamycin A for 2 hours at 30°C in the culture media. For media containing fatty acids, YPAD was supplemented to a final concentration of 1 mM with oleic acid + 1% Tween 80, or 1 mM palmitic acid + 1% Tween 40.

### MDC Assays

Cells were grown overnight at 30°C to saturation in 3 mL of YPAD or SD media. 1 μL of the saturating culture was diluted into 50 mL of fresh media and incubated with shaking for 15-16 hours until the OD600 was between 0.2 and 0.8. 5 mL of the log phase culture was treated with 5 μL DMSO or rapamycin (200 nM final concentration) for 2 hours. Prior to imaging, cells were harvested by centrifugation for 1 minute at 9000 rpm and resuspended in imaging buffer (5% glucose, 10mM HEPES pH 7.6).

For all MDC assays, an n of 3 with 100 cells per n was quantified. In figures, error bars = SEM, p-value as indicated by One Way ANOVA, and Scale Bar = 5 microns.

### Microscopy and Image Analysis

Yeast were directly plated onto a slide at small volumes to allow the formation of a monolayer, and optical z-sections of live yeast cells were acquired with a ZEISS Axio Imager M2 equipped with a ZEISS Axiocam 506 monochromatic camera, 100x oil-immersion objective (plan apochromat, NA 1.4). For super-resolution confocal fluorescence microscopy, a ZEISS LSM800 equipped with an Airyscan detector, 63× oil-immersion objective (plan apochromat, NA 1.4) at room temperature was used. Max projections of individual channels were processed in FIJI. To measure fluorescence intensity in FIJI, 8 x 8 pixel boxes in non-adjusted, single z-sections were measured either covering the MDC or the mitochondrial tubule. To measure diameter of MDCs in FIJI, the line tool was used to draw a line across the diameter of an MDC in non-adjusted, single z-sections where the MDC appeared the largest.

### Statistical Analysis

Prism (GraphPad) was used to perform statistical analysis. The number of replicates, what n represents, and dispersion and precision measures are indicated in the Figure Legends. In general, quantifications show the mean and standard error from three biological replicates with n = 100 cells per experiment. In experiments with data depicted from a single biological replicate, the experiment was repeated with the same results. For lipidomic and metabolomic analysis, MetaboAnalyst was used and graphed using Prism (GraphPad). Input data for each experiment are listed in Supplementary Tables 1, 2, and 3.

### Whole Cell Lysate Preparation and Immunoblotting

Cells were grown as described. 2-10 ODs of cells were lysed in ice-cold 1 mL 0.2 M NaOH/ 0.2% β-mercaptoethanol and incubated on ice for 10 min. 100 μL trichloroacetic acid (TCA) was added to the lysates and incubated on ice for 5 min. Precipitated proteins were harvested via centrifugation at 13,000 rpm for 5 min. Pellets were resuspended in 100 μL 2X SDS sample buffer (0.12M Tris-HCl (pH 6.8), 19% Glycerol, 0.15mM Bromophenol Blue, 3.8% SDS, 0.05% β-mercaptoethanol). 20 μL of 1M Tris base (pH 11) was added and the samples were heated at 75°C for 10 min. Protein samples were loaded on 4-12% SDS-PAGE gels (Bio-Rad) and run at 70-100V. Proteins were semi-dry transferred onto nitrocellulose membrane. Membranes were blocked in PBS-T (137 mM NaCl, 2.7 mM KCl, 10 mM Na_2_HPO4, 1.8 mM KH_2_PO_4_, 0.5% Tween 20) + 5% milk before incubation in the primary antibodies indicated. Membranes were washed 3 times, 10 min each in PBS-T, incubated in secondary antibodies, washed again, and developed with West Atto (Invitrogen). For immunoblot analyses, mouse anti-GFP (1:1,000; Roche), mouse anti-Pgk1 (1:10,000; Invitrogen), and goat-anti-rabbit or donkey-anti-mouse HRP-conjugated secondary (1:5,000, Sigma-Aldrich) were used. Antibody signal was detected with a BioRad Chemidoc MP system. All blots were exported as TIFFs and cropped in Adobe Illustrator.

### Isolation of Yeast Mitochondria

Crudely purified mitochondria were isolated from yeast cells as described in (Schuler et al., 2021). Briefly, yeast were grown overnight in log-phase to an OD600=0.5-1 as described above, then isolated by centrifugation, washed with dH2O and the pellet weight was determined. Cells were then resuspended in 2 mL/g pellet dithiothreitol (DTT) buffer (0.1 M Tris, 10 mM DTT, pH 9.4) and incubated for 20 minutes at 30°C under constant shaking. After re-isolation by centrifugation, DTT treated cells were washed once with zymolyase buffer (1.2 M sorbitol, 20 mM K2HPO4, pH 7.4 with HCl) and cell walls were digested for 30 minutes at 30°C under constant shaking in 7 mL zymolyase buffer per g cell pellet containing 1 mg zymolyase 100T per g cell pellet. After zymolyase digestion, cells were reisolated by centrifugation, washed with zymolyase buffer and lysed by mechanical disruption in 6.5 mL per g pellet homogenization buffer (0.6 M sorbitol, 10 mM Tris pH 7.4, 1 mM ethylenediaminetetraacetate (EDTA) pH 8.0 with KOH, 0.2% BSA, 1 mM phenylmethylsulfonylfluoride) at 4°C. Cell debris were removed from the homogenate twice by centrifugation at 5000 x *g* for 5 min at 4°C and mitochondria were pelleted at 17500 g for 12 min at 4°C. The mitochondrial pellet was resuspended in SEM buffer (250 mM sucrose, 1 mM EDTA pH 8.0 with KOH, 10 mM 3-(N-morpholino)-propansulfonic acid pH 7.2), reisolated by centrifugation at 17500 x *g* for 12 min, resuspended in SEM buffer and mitochondria were shock frozen in liquid nitrogen and stored at −80°C.

### Blue-Native PAGE

Mitochondria were isolated as described and protein complexes were solubilized on ice for 15 min in 1X NativePAGE sample buffer (Thermo Fisher Scientific) with 1% digitonin. Non-solubilized membrane fractions were removed by centrifugation at 20,000 x *g* for 30 min at 4°C. The protein content was determined by a bicinchoninic assay (Thermo Fisher Scientific). 0.25% Coomassie G-250 was added to samples before separation by electrophoresis on a NativePAGE 4-16% Bis-Tris Gel (Thermo Fisher Scientific). Proteins were then transferred to a PVDF membrane (Millipore Sigma) via wet transfer in NuPAGE Transfer Buffer (Thermo Fisher Scientific) at 4°C. Membranes were incubated in 8% acetic acid at RT for 15 min to fix proteins, and then washed in methanol for 5 min to removed background Coomassie G-250. Membranes were blocked in PBS-T + 5% milk before incubation in the primary antibodies indicated. Membranes were washed 3 times, 10 min each in PBS-T, incubated in secondary antibodies, washed again, and developed with West Atto (Invitrogen). For immunoblot analyses, mouse anti-GFP (1:1,000; Roche) and donkey-anti-mouse HRP-conjugated secondary (1:5,000, Sigma-Aldrich) were used. Antibody signal was detected with a BioRad Chemidoc MP system. All blots were exported as TIFFs and cropped in Adobe Illustrator.

### Extraction of Lipids from Yeast Whole-Cell Lysates

Lipids were extracted from yeast as previously described (Xiao et al., 2024). For analysis of whole-cell lysate lipid levels, cells were grown exponentially in the indicated media for 15 h at 30°C to a density of 6–8 × 10^6^ cells/mL. A total of 5 × 10^7^ yeast cells were harvested by centrifugation, washed twice with double-distilled water, and cell pellets were shock-frozen in liquid nitrogen. Extraction of lipids was carried out using a biphasic solvent system of cold methanol, methyl tert-butyl ether (MTBE), and water as described (Matyash et al., 2008) with some modifications. In a randomized sequence, yeast lipids were extracted in bead-mill tubes (glass 0.5 mm; Qiagen) containing a solution of 230 µl MeOH containing internal standards (Cholesterol-d7 [75 μg/mL], and FA 16:0-d31 [28.8 μg/mL] all at 10 μL per sample; Avanti SPLASH LipidoMix) and 250 μL ammonium bicarbonate. Samples were homogenized in one 30-s cycle, transferred to microcentrifuge tubes (polypropylene 1.7 mL; VWR) containing 750 μL MTBE, and rested on ice for 1 h with occasional vortexing. Samples were then centrifuged at 15,000 x *g* for 10 min at 4°C and the upper phases were collected. A 1 mL aliquot of the upper phase of MTBE/MeOH/water (10:3:2.5, vol/vol/vol) was added to the bottom aqueous layer followed by a brief vortex. Samples were then centrifuged at 15,000 x *g* for 10 min at 4°C and the upper phases were combined and evaporated to dryness under speedvac. Lipid extracts were reconstituted in 500 μL of mobile phase B and transferred to a liquid chromatography–mass spectrometry (LC–MS) vial for analysis. Concurrently, a process blank sample was prepared and then a pooled quality control (QC) sample was prepared by taking equal volumes (∼50 μL) from each sample after final resuspension.

### LC-MS Analysis (QTOF)

Lipid extracts were separated on an Acquity UPLC CSH C18 column (2.1 × 100 mm; 1.7 μm) coupled to an Acquity UPLC CSH C18 VanGuard precolumn (5 × 2.1 mm; 1.7 μm) (Waters) maintained at 65°C connected to an Agilent HiP 1290 Sampler, Agilent 1290 Infinity pump, and Agilent 6545 Accurate Mass Q-TOF dual AJS-ESI mass spectrometer (Agilent Technologies). Samples were analyzed in a randomized order in both positive and negative ionization modes in separate experiments acquiring with the scan range m/z 100–1700. For positive mode, the source gas temperature was set to 225°C, with a drying gas flow of 11 liters/min, nebulizer pressure of 40 psig, sheath gas temp of 350°C, and sheath gas flow of 11 l/min. VCap voltage is set at 3500 V, nozzle voltage 500 V, fragmentor at 110 V, skimmer at 85 V, and octopole RF peak at 750 V. For negative mode, the source gas temperature was set to 300°C, with a drying gas flow of 11 l/min, a nebulizer pressure of 30 psig, sheath gas temp of 350°C, and sheath gas flow 11 l/min. VCap voltage was set at 3,500 V, nozzle voltage 75 V, fragmentor at 175 V, skimmer at 75 V, and octopole RF peak at 750 V. Mobile phase A consisted of ACN:H2O (60:40, vol/vol) in 10 mM ammonium formate and 0.1% formic acid, and mobile phase B consisted of IPA:ACN:H2O (90:9:1, vol/vol/vol) in 10 mM ammonium formate and 0.1% formic acid. For negative mode analysis, the modifiers were changed to 10 mM ammonium acetate. The chromatography gradient for both positive and negative modes started at 15% mobile phase B then increased to 30% B over 2.4 min, it then increased to 48% B from 2.4 to 3.0 min, then increased to 82% B from 3 to 13.2 min, then increased to 99% B from 13.2 to 13.8 min where it is held until 16.7 min and then returned to the initial conditions and equilibrated for 5 min. The flow was 0.4 mL/min throughout, with injection volumes of 5 μL for positive and 10 μL negative mode. Tandem mass spectrometry was conducted using iterative exclusion, the same LC gradient at collision energies of 20 and 27.5 V in positive and negative modes, respectively.

### LC-MS Data Processing

For data processing, Agilent MassHunter (MH) Workstation and software packages MH Qualitative and MH Quantitative were used. The pooled QC (*n* = 8) and process blank (*n* = 4) were injected throughout the sample queue to ensure reliability of the acquired lipidomics data. For lipid annotation, accurate mass and MS/MS matching were used with the Agilent Lipid Annotator library and LipidMatch (Koelmel et al., 2017). Results from the positive and negative ionization modes from the Lipid Annotator were merged based on the class of lipid identified. Data exported from MH Quantitative were evaluated using Excel where initial lipid targets are parsed based on the following criteria. Only lipids with relative standard deviations (RSD) <30% in QC samples are used for data analysis. Additionally, only lipids with background AUC counts in process blanks that are <30% of QC are used for data analysis. The parsed Excel data tables are normalized based on the ratio to class-specific internal standards.

Volcano plots were generated in PRISM (GraphPad), based on the MetaboAnalyst Input (Supplementary Tables 1, 2). All phospholipid species were included for PA, PC, PI, PE, PS, PG, and CL.

### Extraction of Whole Cell Metabolites from Yeast

For analysis of whole cell metabolite analysis, yeast cells were grown for 15-17 hours to a density of 0.5-0.9 x 10^7^ cells/mL. 4 x 10^7^ cells/mL were harvested by centrifugation for 3 minutes at 5000g, washed once with water and cell pellets were shock frozen in liquid nitrogen. Whole cell metabolites were extracted from yeast cell pellets as previously described, with slight modifications (Canelas et al., 2009). Briefly, 0.4 μg of the internal standard succinic-d4 acid was added to each sample. Next, 1ml of boiling 75% EtOH was added to each pellet, followed by vortex mixing and incubation at 90°C for 3 min with intermittent vortex mixing. Cell debris were removed by centrifugation at 7000 x *g* for 5 minutes at −10°C. Supernatants were transferred to new tubes and dried *en vacuo*. Process blank samples were made using only extraction solvent and no cell culture pellet.

### GC-MS Analysis for Metabolites

GC-MS analysis was performed with an Agilent 5977b GC-MS MSD-HES fit with an Agilent 7693A automatic liquid sampler. Dried samples were suspended in 40 μL of a 40 mg/mL O-methoxylamine hydrochloride (MOX) (MP Bio #155405) in dry pyridine (EMD Millipore #PX2012-7) and incubated for one hour at 37°C in a sand bath. 25 μL of this solution was added to auto sampler vials.10 μL from the remaining solution for every sample was used to create pooled QC and 25 μL of pooled QC was added to auto sampler vials. 60 μL of N-methyl-N-trimethylsilyltrifluoracetamide (MSTFA with 1% TMCS, Thermo #TS48913) was added automatically via the auto sampler and incubated for 30 minutes at 37°C. After incubation, samples were vortexed and 1 μL of the prepared sample was injected into the gas chromatograph inlet in the split mode with the inlet temperature held at 250°C. A 5:1 split ratio was used for analysis for the majority of metabolites. Any metabolites that saturated the instrument at the 5:1 split were analyzed at a 50:1 split ratio. The gas chromatograph had an initial temperature of 60°C for one minute followed by a 10°C/min ramp to 325°C and a hold time of 10 min. A 30-meter Agilent Zorbax DB-5MS with 10 m Duraguard capillary column was employed for chromatographic separation. Helium was used as the carrier gas at a rate of 1 mL/min.

Data was collected using MassHunter software (Agilent). Metabolites were identified and their peak area was recorded using MassHunter Quant. This data was transferred to an Excel spread sheet (Microsoft, Redmond WA). Metabolite identity was established using a combination of an in-house metabolite library developed using pure purchased standards, the NIST library and the Fiehn library. Values for each metabolite were normalized to the internal standard in each sample, normalized to sum and are displayed as fold change compared to the control sample. Data was analyzed using the in-house ‘MetaboAnalyst’ software tool. The bar graph was generated in PRISM (GraphPad) based on the input data in Supplementary Table 3.

## Abbreviations

OMM: outer mitochondrial membrane
MDC: mitochondrial-derived compartment
UFA: unsaturated fatty acid
TOM: translocase of the outer membrane
TA: tail-anchored
PA: phosphatidic acid
DG: diglyceride
TG: triglyceride

